# Quantitative translation of dog-to-human aging by conserved remodeling of epigenetic networks

**DOI:** 10.1101/829192

**Authors:** Tina Wang, Jianzhu Ma, Andrew N. Hogan, Samson Fong, Katherine Licon, Brian Tsui, Jason F. Kreisberg, Peter D. Adams, Anne-Ruxandra Carvunis, Danika L. Bannasch, Elaine A. Ostrander, Trey Ideker

## Abstract

Mammals progress through similar physiological stages during life, from early development to puberty, aging, and death. Yet, the extent to which this conserved physiology reflects conserved molecular events is unclear. Here, we map common epigenetic changes experienced by mammalian genomes as they age, focusing on evolutionary comparisons of humans to dogs, an emerging model of aging. Using targeted sequencing, we characterize the methylomes of 104 Labrador retrievers spanning a 16 year age range, achieving >150X coverage within mammalian syntenic blocks. Comparison with human methylomes reveals a nonlinear relationship which translates dog to human years, aligns the timing of major physiological milestones between the two species, and extends to mice. Conserved changes center on specific developmental gene networks which are sufficient to capture the effects of anti-aging interventions in multiple mammals. These results establish methylation not only as a diagnostic age readout but as a cross-species translator of physiological aging milestones.

## INTRODUCTION

A key molecular hallmark of vertebrate aging is remodeling of the DNA methylome, e.g. the pattern of epigenetic modifications whereby methyl groups are present at some cytosine-guanine dinucleotides (CpGs) but absent from others (Field et al. 2018). Although the several species examined thus far display some degree of epigenetic remodeling with age, several questions remain. First, the rate of methylation change appears to depend on the maximal lifespan of the species (i.e. the relative time at end of life) (Maegawa et al. 2017; Lowe et al. 2018). However, whether maximal lifespan is the sole factor in determining how the methylome progresses with age, or if the methylome is entrained by additional milestones in vertebrate development and aging, remains to be investigated. Second, it is unclear if the major epigenetic changes with age involve the same, different, or random CpG sites in distinct species. While DNA encoding a highly conserved ribosomal RNA family shows increasing methylation over time at the same conserved CpG sites in many species (M. Wang and Lemos 2019), methyl-CpGs associated with age in humans by so-called “epigenetic clocks” (Hannum et al. 2013; S. Horvath 2013; T. Wang et al. 2017; Petkovich et al. 2017; Thompson et al. 2017; Stubbs et al. 2017) have not been strongly predictive of age in other mammals (Petkovich et al. 2017; Stubbs et al. 2017). This result is inconclusive, however, since epigenetic clocks typically use fewer than 500 CpG sites for age prediction out of the potential millions of CpG sites that make up the mammalian methylome.

Thus far, an important impediment to answering these and other questions has been our inability to characterize conserved epigenetic features in different species. Commonly-used techniques such as whole-genome bisulfite sequencing (WGBS) and reduced representation bisulfite sequencing (RRBS) measure large numbers of random CpGs, but such CpGs are often not present in regions conserved across species. Thus how, where, and to what extent epigenetic remodeling reflects a conserved process in aging is still largely unaddressed.

Domestic dogs provide a unique opportunity to investigate the above questions (Kaeberlein, Creevy, and Promislow 2016; Gilmore and Greer 2015). Dogs have been selectively bred by humans for occupation and aesthetics (Ostrander et al. 2017), resulting in over 450 distinct whose members share morphologic and behavior traits. Most breeds derive from small numbers or popular sires, leading to strong phenotypic and genetic homogeneity within breeds (Dreger et al. 2016).

Although humans and dogs diverged early during mammalian evolution (Vonholdt et al. 2010), dogs share nearly all aspects of their environments with humans. These include, critically for these studies, similar levels of health observation and health care intervention (Kaeberlein, Creevy, and Promislow 2016; Gilmore and Greer 2015). While average lifespan differs dramatically across breeds, within some breeds such as the Labrador retriever there are extensive variations in lifespan. In these cases, use of a single breed offers the advantage of strong genome homogeneity, increasing the probability of identifying genetic factors associated with any complex trait (Davis and Ostrander 2014), including aging. Despite extensive differences, all dogs exhibit a similar developmental, physiological and pathological trajectory (Kaeberlein, Creevy, and Promislow 2016; Gilmore and Greer 2015), but they accomplish this progression in many fewer years than do humans, generally less than 20. Finally, epigenetic clocks have been demonstrated in dogs (Thompson et al. 2017) thus increasing their utility as a system for studies of age-related epigenetic remodeling in humans (Hannum et al. 2013; S. Horvath 2013).

## RESULTS

To enable high quality evolutionary comparisons of dog methylomes with other mammals, we performed targeted-bisulfite sequencing to systematically characterize CpGs in regions of the dog genome that are syntenic with those measured by Illumina human methylome arrays. Since Illumina arrays have been used to characterize epigenetic aging in many human studies (Hannum et al. 2013; S. Horvath 2013; Alisch et al. 2012), our goal was to create a high-quality panel of dog methylomes with substantial coverage of CpGs noted in prior human datasets (**Figure 1A**). Our strategy, henceforth called *Synteny Bisulfite Sequencing* (SyBS), was designed to capture approximately 90,000 CpGs of the approximately 232,000 conserved CpGs on the Illumina array (**Methods**).

**Figure 1:**
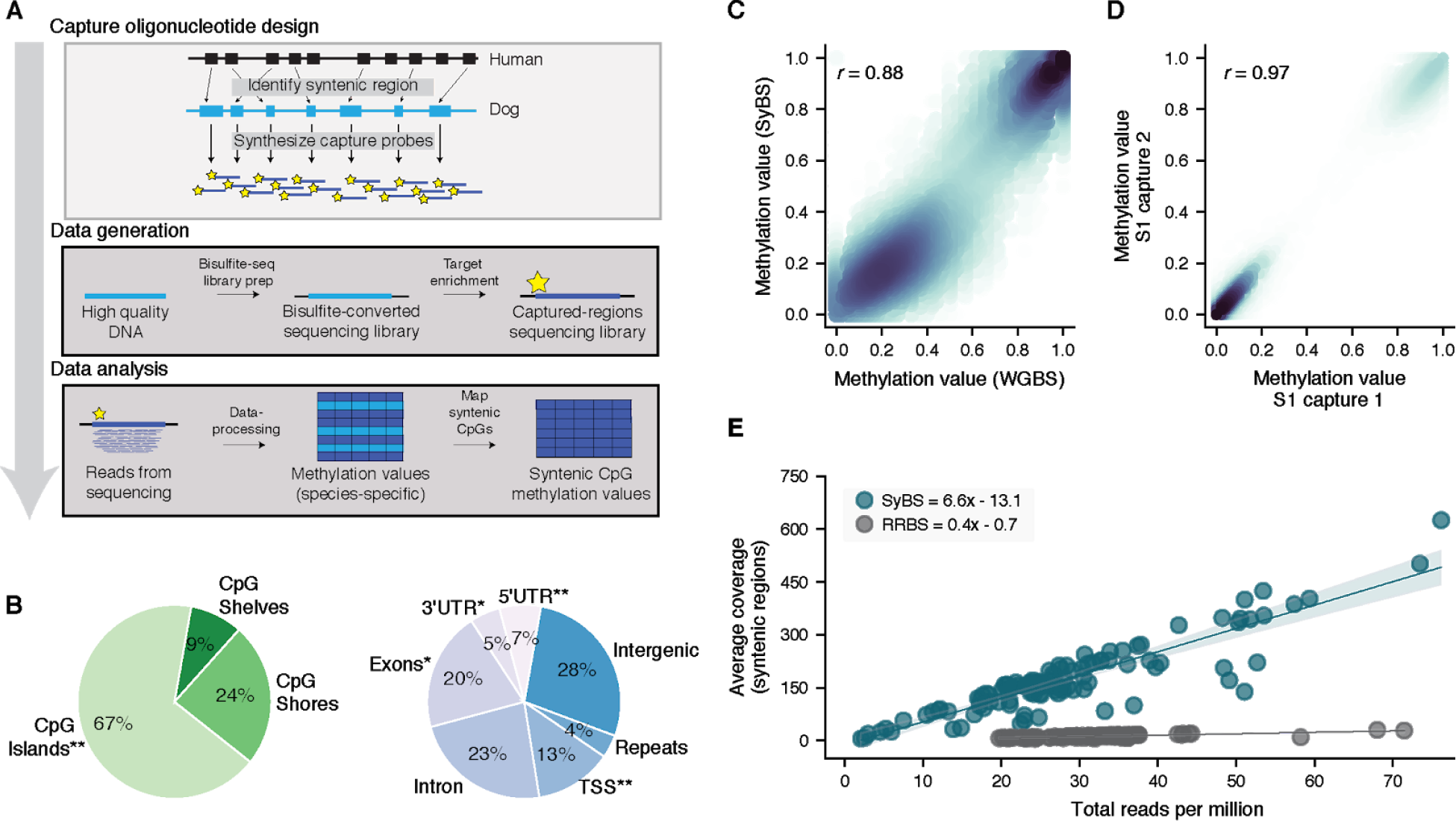
Interrogating mammalian methylomes by syntenic bisulfite sequencing (SyBS). (**A**) Strategy used to profile and compare CpG methylation states within blocks of synteny in the mammalian genome. Capture oligonucleotide design: Regions of DNA (blue blocks) characterized by the Illumina 450K methylation array in humans are mapped to their syntenic region in dogs using whole-genome alignments between the two species. These regions are used to design oligonucleotides (yellow stars) for capture and enrichment of DNA in the second species. Data generation: A sequencing library is constructed from high quality DNA and bisulfite converted, analogously to whole-genome bisulfite sequencing. Syntenic sequences are captured, sequenced, and aligned to the mammalian genome under study. CpG methylation values are called then filtered to select those conserved with humans for further analysis. For more details see **Methods**. (**B**) Pie charts showing representation of targeted genomic regions. Regions exhibiting significant enrichment (p < 10^−10^) are indicated using asterisks with * indicating odds ratio > 2.5 and ** odds ratio > 4. (**C**) Ten dog methylomes were sequenced twice, either with enrichment for syntenic regions (SyBS hybridization) or without enrichment (whole-genome bisulfite sequencing, WGBS). Methylation values (per CpG site per animal) are shown for SyBS (y-axis) versus WGBS (x-axis). Sites were considered if they were covered by >5 reads with both SyBS and WGBS. (**D**) Concordance of SyBS values for one canine DNA sample (S1), for which two independent captures were performed. In panels (C) and (D) the color captures the density of observations at each point (darker colors represent higher densities), and the *r*-value is the Pearson correlation. (**E**) Average coverage of syntenic segments versus total reads in millions, contrasting SyBS with Reduced Representation Bisulfite Sequencing (RRBS).

We applied SyBS to characterize the methylomes of 104 dogs, primarily consisting of Labrador retrievers and representing the entire lifespan, from 0.1 to 16 years at the time of blood draw (**Figure S1A**, **Table S1**). Libraries were sequenced to an average depth of 163X, with nine dogs removed due to lack of coverage. Captured CpG sites spanned the genome and were enriched for regions including exons, transcription start sites and CpG islands (odds ratio > 2.5 and p < 10^−10^ by Fisher’s exact test, **Figure 1B**). The methylation values associated with captured CpGs were similar to those obtained using whole-genome bisulfite sequencing (Pearson correlation *r* > 0.85, **Figure 1C**) and showed excellent replication across independent captures from the same samples (*r* > 0.95, **Figure 1D**, **Figures S1B-F**). As expected, SyBS achieved approximately 13-times higher coverage of syntenic regions compared to non-targeted reduced representation bisulfite methods (**Figure 1E**). For comparison, we obtained previously published methylation profiles from the blood of 320 human individuals aged one to 103 years at the time of sample isolation (Alisch et al. 2012; Hannum et al. 2013). Based on these data, we identified 54,469 well-profiled CpGs in both species, thus enabling systematic evolutionary studies of epigenetic changes during life (**Methods**).

We observed the highest methylome similarities (Pearson correlation, **Methods**) when pairing young dogs with young humans, or aged dogs with older humans. In contrast, the lowest similarities were obtained when pairing young dogs with old humans or *vice versa* (**Figure 2A**). The relationship between methylome similarity and age was lost upon permutation (FDR < 0.01; **Figure S2A**), indicating that a conserved set of CpG sites are affected during aging in the two mammalian species. Notably, this signal was sufficiently strong to arise in an unbiased methylome-wide analysis without subselection of markers. This result suggests that the conserved methylation changes with age in ribosomal RNA genes, noted previously (M. Wang and Lemos 2019), extend more generally to the greater mammalian methylome. It contrasts with previous observations using epigenetic clocks, which did not find strong conservation across species, likely because these clocks are restricted to 80-300 CpGs selected for optimal age prediction in humans independent of other species (Stubbs et al. 2017).

**Figure 2:**
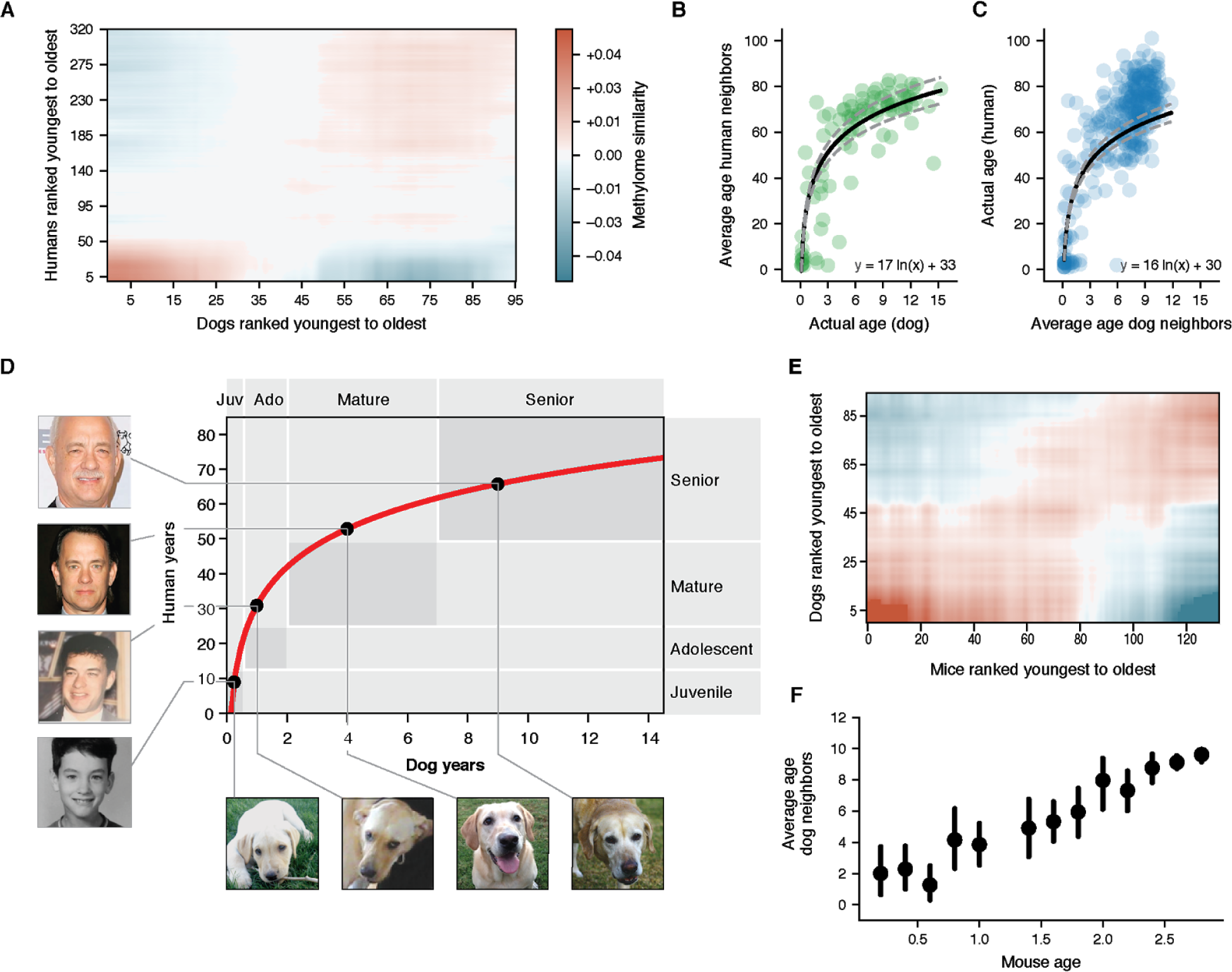
A non-linear transformation from dog to human age. (**A**) Dog-human methylome similarities (Pearson correlation, blue-red color range) are shown with dogs and humans ranked from youngest to oldest. Data are lightly smoothed in both dimensions using Gaussian interpolation in matplotlib. (**B**) The age of each dog methylome (x-axis) is plotted against the average age of the five nearest human methylomes (y-axis). (**C**) Reciprocal plot in which the age of each human methylome (y-axis) is plotted against the average age of the five nearest dog methylomes (x-axis). (**D**) Logarithmic function for epigenetic translation from dog age (x-axis) to human age (y-axis). Outlined boxes indicate the approximate age ranges of major life stages as documented qualitatively based on common aging physiology. Juvenile refers to the period after infancy and before puberty, 2-6 mos. in dogs, 1-12 yrs. in humans; Adolescent refers to the period from puberty to completion of growth, 6 mos. to 2 yrs. in dogs, approximately 12-25 yrs. in humans; Mature refers to the period from 2-7 yrs. in dogs and 25-50 yrs. in humans; Senior refers to the subsequent period until life expectancy, 12 yrs. in dogs, 70 yrs. in humans. Dog life stages are based on veterinary guides and mortality data for dogs (Bartges et al. 2012; Inoue et al. 2015; Fleming, Creevy, and Promislow 2011). Human life stages are based on literature summarizing life cycle and lifetime expectancy (Bogin and Smith 1996; Cia 2013; Arias, Heron, and Xu 2017). Black dots on the curve connect to images of the same yellow Labrador taken at four different ages (courtesy of Sabrina and Michael Mojica, with permission) and to images of a representative human at the equivalent life stages in human years (photos of Tom Hanks drawn from a public machine-learning image repository (Chen, Chen, and Hsu 2015)). (**E**) Mouse-dog methylome similarities shown as in panel (A). (**F**) Data from panel (E) are summarized by sorting mice according to 0.2 year bins (x-axis) and, for each mouse, plotting the average age of the 5 nearest dogs by methylome similarity (y-axis). Points illustrate the mean of each bin and bars represent the 95% confidence interval obtained from bootstrapping.

We next investigated whether the conserved methylation changes in dogs and humans showed a constant rate of change with age, with the rate constant depending on the lifespan of each species as suggested by previous studies (Lowe et al. 2018; Maegawa et al. 2017), or whether there was evidence for a more complex trajectory. For this purpose we assigned the age of each dog to the average age of its nearest *k* humans by methylome-wide similarity (**Methods**). This analysis revealed a monotonic, time-resolved, nonlinear relationship between dog and human age (**Figure 2B** for *k* = 5, **Figures S2B-G** for other *k*). Similar results were obtained in a reciprocal analysis assigning each human to its nearest dogs (**Figure 2C**), allowing us to combine the reciprocal analyses to generate the single function: *human_age* = 16 *ln*(*dog_age*) + 31 (**Figure 2D**).

We found that this function showed strong agreement between the approximate times at which dogs and humans experience common physiological milestones during both development and lifetime aging, i.e. infant, juvenile, adolescent, mature, senior (Lebeau 1953; Bogin and Smith 1996; Bartges et al. 2012) (**Figure 2D**). The observed agreement between epigenetics and physiology was particularly close for infant, juvenile and senior stages. For instance, the epigenome translated approximately 8 weeks in dogs (0.15 years) to approximately nine months in humans (0.78 years), corresponding to the infant stage when deciduous teeth develop in both puppies and babies (Bogin and Smith 1996; Bartges et al. 2012). In seniors, the expected lifespan of Labrador retrievers, 12 years, correctly translated to the worldwide lifetime expectancy of humans, 70 years (Cia 2013; Fleming, Creevy, and Promislow 2011). For adolescent and mature stages, the correspondence was more approximate, with the epigenome showing faster changes for dogs, relative to humans, than expected by physiological tables (Inoue et al. 2015; Arias, Heron, and Xu 2017) (**Figure 2D**). Thus, the canine epigenome progresses through a series of conserved biological states which align with major physiological changes in humans, occurring in the same sequence but at different chronological timepoints during each species’ lifespan.

A conserved nonlinear and weaker, epigenetic progression was also observed by comparing the dog methylomes to those from the 133 mice (Petkovich et al. 2017) (**Figures 2E-F**). This weaker effect may be due to the limited number of mice sampled during the developmental period (**Table S3**). Nevertheless, the ability to translate age among these three diverse mammals indicates that shared physiology may yield conserved molecular transitions in epigenome remodeling with age.

To determine whether the conserved changes were concentrated within particular genes or gene functions, we examined CpG methylation states near 7,942 genes for which orthologs were present in all three species (dogs, humans, and mice, **Methods**). This analysis identified 394 genes for which methylation values showed conserved time-dependent behavior across species (empirical p < 0.05, **Figure S3, Table S2**). To understand the underlying gene functions we mapped them onto the Parsimonious Composite Network (PCNet), a database of approximately 2×10^6^ molecular interactions capturing physical and functional relationships among genes and gene products, in which each interaction has support from multiple sources (Huang et al. 2018). The genes clustered into five highly interconnected network modules (**Figure 3**), nearly all of which were enriched for developmental functions. Four modules predominantly increased methylation with age (FDR < 0.05), and included modules associated with synapse assembly (18 genes); neuroepithelial cell differentiation (five genes) and two modules associated with anatomical patterning (117 and 69 genes). The four modules were enriched for polycomb repressor targets, which are predominantly silenced in adult tissues (Xie et al. 2013). A fifth module was enriched in leukocyte differentiation and nucleic-acid metabolism (144 genes) and demonstrated decreasing methylation with age. We also noted that orthologs from all five modules were among the most highly conserved in DNA sequence in the mammalian genome, even accounting for high sequence conservation of developmental genes, in general (**Figure S4**). A further indication of the importance of developmental gene modules was observed when calculating dog-human methylome similarity using CpGs at developmental genes only, versus a comparison using all CpGs except those at developmental genes (**Figure S5**, **Methods**). Thus, CpGs near developmental genes were both necessary and sufficient to recapitulate the cross-species alignments of age observed earlier (**Figure 2**). The enrichment of age-related methylation increases in developmental genes has been previously observed in humans (Rakyan et al. 2010) and mice (Maegawa et al. 2010). Our findings further extend such observations by showing they are predominant drivers of the ability to align mammalian epigenomes, and in specifying more precisely where in developmental gene modules such changes occur.

**Figure 3:**
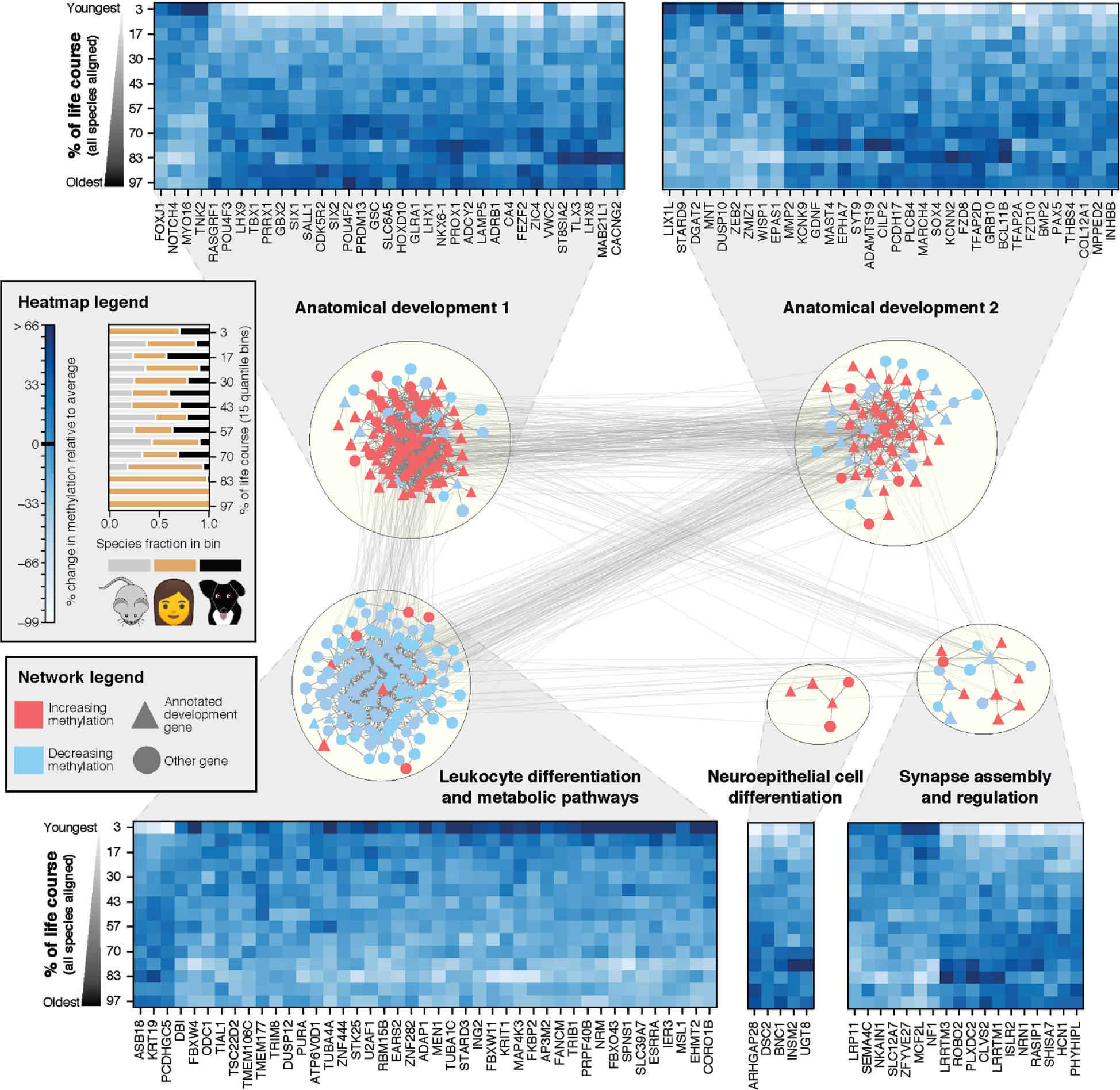
Conserved lifetime methylation changes aggregate in functional networks. Genes exhibiting conserved age-related methylation behavior were mapped onto a composite molecular interaction network which was subsequently clustered to reveal five major modules, labeled according to enriched Gene Ontology functions (**Methods**). Colors represent the conserved direction of change with age, with red representing genes that increase in methylation with age, and blue representing genes that decrease in methylation with age. Heatmaps show the conserved methylation patterns of a random subset of genes in each module. Columns represent distinct orthologs, while rows represent the average values of all species ranked according to their age in human years and divided into 15 age bins (quantiles). Values are normalized according to the mean and standard deviation of methylation for each ortholog. The fractional species composition of each bin is visualized in the legend.

If the methylation pattern associated with developmental modules tracks the physiological progression of the organism, and not just chronological time, we hypothesized that this methylation pattern will respond to interventions that slow or delay aging, such as anti-aging treatments. In mice, calorie restriction and dwarfism have been associated with increased lifespan relative to controls (Petkovich et al. 2017). We therefore examined whether such effects were observed within conserved development gene modules. For this purpose we used the methylation patterns of developmental modules to construct an epigenetic clock, a regularized linear regression model that measures age from CpG methylation values (Hannum et al. 2013; S. Horvath 2013; T. Wang et al. 2017; Petkovich et al. 2017; Thompson et al. 2017; Stubbs et al. 2017) (439 CpGs total, **Figure 4A** and **Methods**). By training in dogs or alternatively mice, this “development clock” could be used to translate a methylation profile to either dog or mouse years (**Figures 4B-C**). Implicit in this is the translation of a dog methylome to its equivalent mouse age or *vice versa* (**Figures 4D-E**). We observed that epigenetic ages measured by the clock were more consistent with the actual ages of animals than were clock measurements formulated from the rest of the methylome (**Figures 4D-E**, controlling for number of CpGs). When applying the development clock to mice treated with lifespan-extending interventions, the epigenetic ages were 30% less, on average, than those of control mice (p < 10^−6^, **Figure 4F**). These same results were observed when using the development clock trained in dogs to predict mouse age (**Figure 4G**). Together, these results demonstrate that the methylation states of developmental gene modules track the physiological effects of aging and aging interventions in multiple mammalian species.

**Figure 4:**
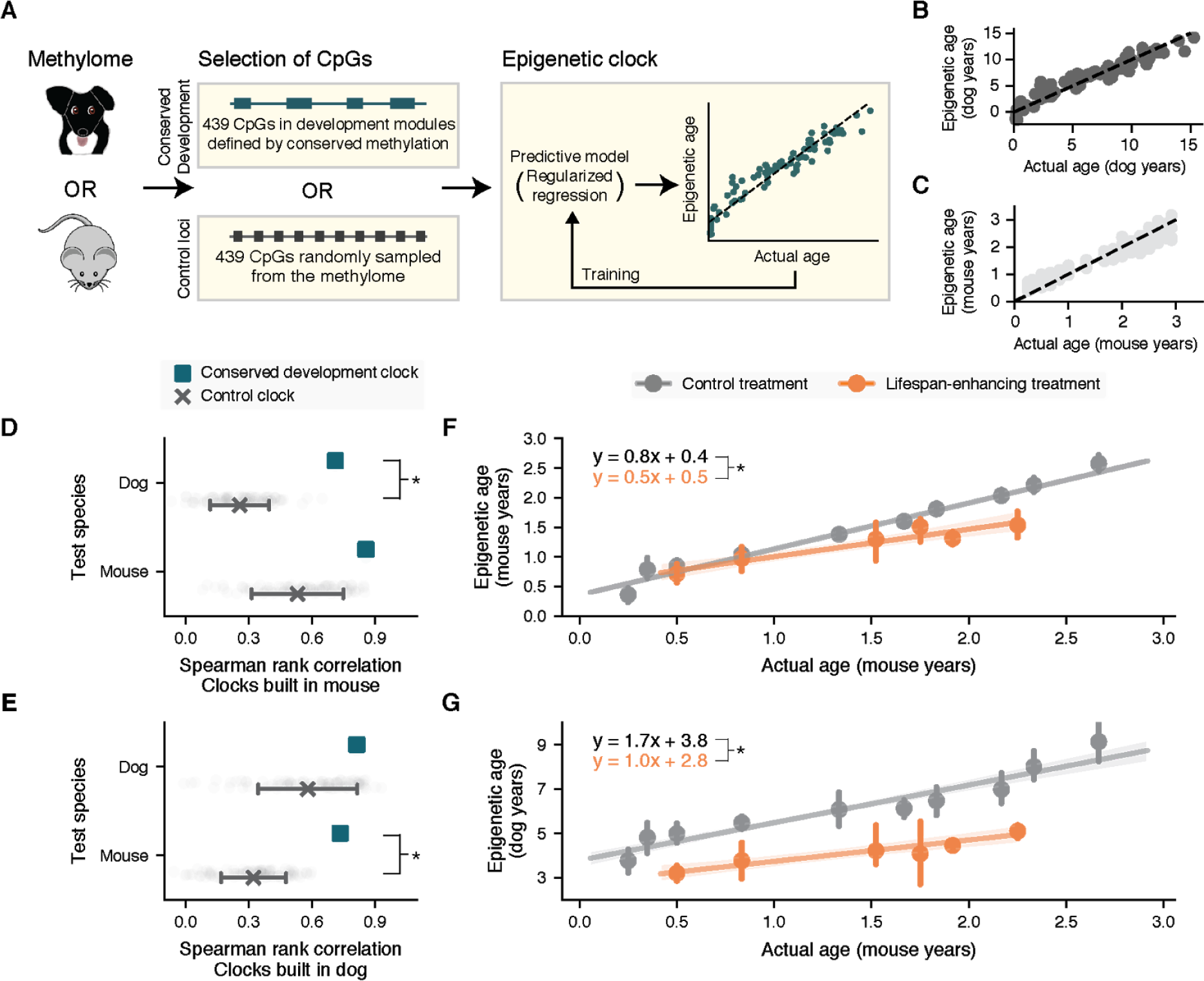
A conserved development clock measuring both age and biological aging. (**A**) Schematic of epigenetic clock construction. The model computes the sum of methylation values from either dogs or mice, using CpGs from developmental gene modules (conserved development clock) or from random samples of the same number of CpG sites (control clock). The relative weight of each CpG is trained to predict the chronological age of each input sample using regularized regression (ElasticNet, **Methods**). Training performance in dogs (**B**) or mice is shown. (**D-E**) Spearman correlation between epigenetic and actual ages (x-axis) when training clocks in mice (**D**) or dogs (**E**), for the test species indicated (y-axis, top dog, bottom mouse). Green and gray points show the performance of the conserved development clock or 100 random control clocks, respectively (mean ± 95% confidence interval). * denotes p < 0.05 in all panels. (**F**) The conserved development clock distinguishes the effects of lifespan-enhancing treatments (orange) from control treatments (gray). For each treatment, mouse epigenetic ages are measured (y-axis, conserved development clock trained in mice) and plotted against actual mouse ages binned in 10 age quantiles (x-axis). Mean ± 95% confidence intervals shown for each bin. (**G**) As for panel (F) but training the conserved development clock using data for dogs. For each treatment (orange lifespan-enhancing, gray control), epigenetic ages of each mouse are measured and plotted against actual mouse ages binned in 10 age quantiles (x-axis). * denotes p < 0.05 in all panels.

## DISCUSSION

By using targeted oligonucleotide capture (SyBS), we have produced epigenomes of high quality for comparative studies between dogs and other mammals. Analysis of these data shows that multiple mammalian species experience conserved methylation changes during aging, and that the scope of these changes is methylome-wide. Second, the trajectory of epigenetic changes followed by one species as it ages is not necessarily the same as that followed by another. In particular, dog methylomes remodel very rapidly in early life compared to the methylomes of their human counterparts. Our further analysis in this regard (**Figure 2D**) suggests that the rate of epigenetic remodeling is not only determined by the lifespan of a species but also by the timing of key physiological milestones. Previously, CpG methylation states in humans were proposed to exhibit a non-linear trajectory over time, with non-constant rates of change (S. Horvath 2013). The analysis here demonstrates a different point, that methylation changes in one species can be non-linear with respect to another.

We observe that epigenetic changes during aging center on highly conserved modules of developmental genes, in which methylation generally increases with age. Although the enrichment of developmental pathways has been generally observed in mammals previously (Field et al. 2018; Ciccarone et al. 2018; Steve Horvath and Raj 2018), our findings show that specific changes at these loci translate the effects of age and anti-aging interventions across species. While the biology of aging has historically been considered as separate from that of development (Miller and Nadon 2000; Kowald and Kirkwood 2016), their strong association, demonstrated here, supports a model in which at least some aspects of aging are a continuation of development rather than a distinct process.

Limitations and extensions of our findings are as follows. First, we have used DNA isolated from whole blood, for which age-dependent shifts in leukocyte populations have been described (Jaffe and Irizarry 2014). In particular, previous studies have found that CD4+ T cells, CD8+ T cells and B cells decline with age. Although it is possible that such conserved shifts may influence our findings, such decline occurs in both dogs (Greeley et al. 2001) and humans (Jaffe and Irizarry 2014). Second, our study has focused exclusively on Labrador retrievers, a popular and heterogenous breed for which we could collect large numbers of unrelated dogs in order to control for population structure. Distinct breeds exhibiting widely varying lifespans (Kaeberlein, Creevy, and Promislow 2016; Gilmore and Greer 2015) could yield different epigenetic age translation functions.

Further efforts to characterize epigenetic changes across species may also help to address broader questions. For example, does the timing of epigenetic changes early in life influence the overall lifespan of a species, or of an individual within that species? Does modulating the timing of developmental events affect lifespan? Again, comparisons of species or sub-species that experience developmental milestones at similar times but with different lifespans (such as distinct dog breeds) may help address these questions, providing critical and complementary data to inform ongoing cross-species aging studies (Kaeberlein, Creevy, and Promislow 2016) including clinical trials of aging interventions (Urfer et al. 2017).

Finally, our study has demonstrated that the methylome can be used to quantitatively translate the age-related physiology experienced by one organism (i.e. a model species like dog) to the age at which physiology in a second organism is most similar (i.e. a second model or humans, **Figure 2**). These results create the opportunity to use the methylome not only as a diagnostic readout of age (as per the epigenetic clock) but for cross-species translation of physiological state. Such translation may provide a compelling tool in the quest to understand aging and identify interventions for maximizing healthy lifespan.

## Supporting information

Table S2: Genes exhibiting conserved time-dependent behavior

Table S3: Mice sample description

## METHODS

### Annotations

Reference genomes were downloaded from Ensembl for dog (CanFam3.1), mouse (mm10) and human (hg19). Ensembl Biomart version 91 was used for gene, 3’UTR and 5’UTR annotations (Yates et al. 2016). CpG islands, repeat annotations, and chain files were downloaded from the UCSC Genome Browser (Rosenbloom et al. 2015). CpG shores were designated as regions 2 kilobases (kb) outside each CpG island, and CpG shelves were designated as regions 2kb outside of CpG shores. Promoters were designated as regions 2kb upstream and 100 basepairs (bp) downstream of the transcription start sites (TSS) based on gene annotations from Ensembl (Yates et al. 2016). Whole genes were divided into exonic and intronic sequences. Intergenic regions were then defined as the remaining regions of the genome after subtracting all other annotated regions. Definitions of one-to-one orthologs were downloaded from Ensembl Compara (Vilella et al. 2009) for dogs, humans and mice.

### Public datasets

The following datasets were obtained from Gene Expression Omnibus (GEO) or Sequence Read Archives (SRA) [number of individuals included in study in brackets]:

- GSE80672 (Petkovich et al. 2017): Methylomes from postnatal mice. Blood, Reduced Representation Bisulfite Sequencing (RRBS) method. [133]
- GSE36054 (Alisch et al. 2012): Methylomes from human children. Blood, Infinium 450K array. [35]
- GSE40279 (Hannum et al. 2013): Methylomes from human adults. Blood, Infinium 450K array. [285]
- SRP065319 (Thompson et al. 2017): Methylomes from dogs and wolves. Blood, RRBS method. [92]

### Canine samples

Information on each dog sample used, including age, breed, and source, is given in **Table S1**, with the age distribution also provided in **Figure S1A**. For samples sourced from NHGRI, domestic dogs were collected with owners’ signed consents in accordance with standard protocols approved by the NHGRI IACUC committee. Samples were collected at canine-centric events such as dog shows. Alternatively, owners were supplied with a mail-in kit which included instructions, tubes for blood draws and a general information sheet requesting the AKC number (when available), pedigree and date of birth. Blood draws were performed by licensed veterinarians or veterinary technicians. For samples sourced from UC Davis, blood was collected from privately owned dogs through the William R. Pritchard Veterinary Medical Teaching Hospital. Owners specified the breed of each dog. Standard collection protocols were reviewed and approved by the UC Davis IACUC. DNA was extracted either using the Puregene kit (Qiagen) or using the cell lysis protocol described by (Bell, Karam, and Rutter 1981), followed by phenol/chloroform extraction with phase separation in 15 mL phase-lock tubes (5-Prime, Inc. Gaithersburg, MD, USA).

### SyBS target selection

The strategy for syntenic bisulfite sequencing was to base our Illumina Human 450K probe locations were extended 50bp with respect to the strand of each probe. The resulting locations were mapped to the dog genome using liftOver (Rosenbloom et al. 2015) using default parameters. After excluding regions that mapped to sex, mitochondrial and unplaced contigs in the dog genome, we identified approximately 230,000 probes that were syntenic between human and dogs. Hybridization probes were generated to target these regions using the Roche SeqCap-Epi platform. This process produced an 18.8 megabase sequencing library in dogs, containing approximately 90,000 CpGs that were also profiled by the Illumina 450K array in humans.

### SyBS library preparation and sequencing

We followed the protocol specified by the Roche SeqCap-Epi platform. Briefly, approximately 500ng of lambda phage DNA (bisulfite-conversion control) was added to 1ug of dog DNA, then sheared to an average of 175bp (Covaris). Sheared DNA was end-repaired, A-tailed and ligated to barcoded adapters. Adapter-ligated libraries were subjected to bisulfite treatment (Zymo EZ DNA methylation lightning kit) following manufacturer’s instructions. Bisulfite-treated libraries were cleaned and amplified using 25 cycles of PCR with a uracil-tolerant enzyme (Kapa). Libraries were pooled equimolarly into 4-plex or 6-plex hybridization capture reactions to a total of 1ug per reaction. Captured product was PCR amplified (10 cycles). Hybridizations were pooled before sequencing and split among 10 lanes on an Illumina HiSeq 4000 in 2×150bp cycles.

### SyBS data analysis

Reads obtained from sequencing were demultiplexed and their quality was verified using FastQC (Andrews and Others 2010). Reads were trimmed using TrimGalore (Krueger 2015) (4bp) then aligned to a bisulfite-converted dog genome (CanFam3.1) using Bismark (v0.14.3) (Krueger and Andrews 2011), which produced alignments with Bowtie2 (v2.1.0) (Langmead et al. 2009) with parameters “-score_min L,0,−0.2”. Methylation values for CpG sites were determined using MethylDackel (v0.2.1) (Ryan 2017). Custom Python scripts using BEDtools (v2.25.0) (Quinlan and Hall 2010) were used to determine on-target reads. Optical PCR duplicates were determined using Picard tools (v1.141) (Tools 2015) and removed using Samtools (v0.1.18) (Li et al. 2009). Coverage of syntenic regions was determined using the number of unique on-target reads that were orthologous to humans, divided by the expected sequencing space. Only CpG sites that were on-target, covered by at least five reads and present across 90% of samples were selected for further analysis. Samples missing more than 30% of CpGs were removed from further analyses resulting in the removal of nine dogs. Missing data for selected CpGs were imputed by performing k-nearest neighbors (*k* = 10) using fancyimpute in Python. To assess the concordance of methylation values obtained using SyBS with conventional approaches, we also sequenced 10 of the same dogs using whole-genome bisulfite sequencing (libraries prior to enrichment with SyBS probes). Reads were processed and aligned with the canine genome as described above. We saw an average Pearson correlation of *r* = 0.85 among these 10 samples (range 0.75 - 0.97) (**Figure 1C**). We also performed independent replicate hybridizations for 6 samples. We saw an average *r* = 0.97 (range: 0.96 - 0.98) for these technical replicates (**Figure 1D, Figure S1B-F**). We verified that lambda phage DNA exhibited complete conversion (>99.5%). We tested the significance of the enrichment of our captured sequences and genome region annotation using the LOLA package (Sheffield and Bock 2016) in R (version 3.5.1) (R Core Team 2018). Enrichment tests are performed using Fisher’s exact tests, with the possible ‘universe’ defined by restriction digestion fragmentation of autosomes in the canine reference genome.

### Public RRBS data processing

For data generated using Reduced Representation Bisulfite Sequencing (RRBS), methods for alignment and CpG selection were identical to those described above. Since RRBS fragments are generated using restriction enzymes with specific recognition sites, optical PCR duplicates could not be removed and on-target CpGs were not determined. For evolutionary comparative analysis, we included 133 control mice aged between 3 months to 2.5 years (Petkovich et al. 2017). To compare the coverage of syntenic regions between SyBS and non-targeted bisulfite technology, we used a RRBS study in dogs and wolves (Thompson et al. 2017) (**Figure 1E**).

### Human methylation array data processing

Illumina Infinium 450K methylome array data were quantile normalized using Minfi (Aryee et al. 2014) and missing values were imputed using the Impute package (Hastie et al. 2011) in R. These values were adjusted for cell counts as previously described (Gross et al. 2016). To enable comparisons across different methylation array studies, we implemented beta-mixture quantile dilation (BMIQ) (Teschendorff et al. 2013; S. Horvath 2013) and used the median of the (Hannum et al. 2013) dataset as the gold standard. To mitigate residual batch effects, we selected human samples that clustered closely in the first two principal components using scikit-learn v0.19.2 (Pedregosa et al. 2011) and verified that such filtering had little effect on the distribution of ages. We also removed samples for which more than 10% of probes were not adequately detected. This procedure resulted in methylome profiles for 320 humans that could be compared to the SyBS-generated dog methylomes.

### Determining orthologous CpGs

Human Illumina 450K methylation array CpGs were extended by 50bp with respect to the strand using BEDtools and mapped to the target genome (mouse or dog) using liftOver with “-minMatch=0.5”. We verified that the coordinate alignment obtained using 50bp was identical to that obtained using the exact coordinate (1bp) at “-minMatch=0.95”. This procedure allowed us to determine an exact orthologous region for each human CpG and each dog CpG. When multiple dog CpGs were assigned to one human CpG probe region, we took the average methylation value of the aligned CpGs in dogs. This procedure resulted in 54,469 dog-human orthologous CpGs for further analysis. To mitigate batch effects specific to sequencing and/or array platforms, we normalized the sequencing methylation values using BMIQ and performed quantile normalization using the preprocessCore package in R (normalize.quantiles.use.target function) (Bolstad 2013).

For dog-to-mouse comparisons, CpGs that were separated by 1bp were merged into one region using BEDtools. Each region was then extended by 50bp. The resulting region files were aligned to the target genome using liftOver “−minMatch=0.5”. Only regions that were concordant between the two alignments (*i.e.*, dog to mouse or mouse to dog) were selected for further analysis. CpGs that were assigned to the same aligned regions were averaged to generate 9,404 bins, consisting of 87,915 CpGs from dogs.

### Computing dog-human pairwise methylome similarity

Methylation values of orthologous CpGs were normalized by subtracting the mean and dividing by the standard deviation over individuals (i.e. z-transformed, separately for each species). The resulting z-values represent the tendency to decrease or increase relative to the mean of each CpG within a species. Using these values we calculated the pairwise Pearson correlation between the methylomes of each dog-human pair. Correlation was computed across all orthologous CpG values using the SciPy Python package (Jones et al. 2015), forming a 95 × 320 (dog × human) methylome similarity matrix (**Figure 2**). We also created a coarsened version of this matrix, in which the pairwise similarities were averaged over two-year age windows in both species, forming an 8 × 51 (dog × human) methylome similarity matrix (*MSA*, **Figure S2A**).

Given this matrix, we evaluated the significance of association between age and methylome similarity using permutations. Specifically, we generated the following two-by-two contingency table:

| | Ages more different<br>$AD(i,j) > \overline{AD}$ | Ages more similar<br>$AD(i,j) \leq \overline{AD}$ |
| --- | --- | --- |
| Methylomes more different<br>$MSA(i,j) \leq \overline{MS}$ | $Count_1$ | $Count_3$ |
| Methylomes more similar<br>$MSA(i,j) > \overline{MS}$ | $Count_2$ | $Count_4$ |

where *MSA* is the methylome similarity matrix, *AD*(*i*, *j*) is the age difference computed as | *Age bin*_*dog*_ − *Age bin*_*human*_ | and *Count* is the number of occurrences (cells within the *MS* similarity matrix) for which the table row and column conditions are met. Using these counts, we calculated the p-value using the one tailed Fisher’s exact test and compared this p-value to that obtained when permuting the membership of dogs and humans in two-year age bins across 1000 permutations (**Figure S2A**).

### k-nearest neighbors analysis

To achieve a robust assignment of reciprocal nearest neighbors, we used a strategy inspired by Context Likelihood of Relatedness (Madar et al. 2010). Specifically, we z-normalized the *MS* methylome similarity matrix to form *MSZ*, as follows:

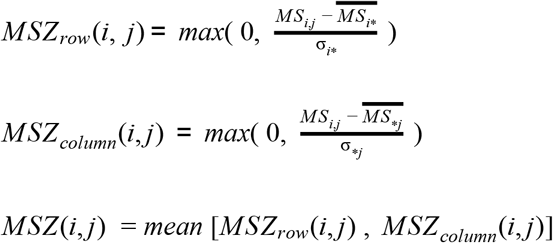

*k*-nearest neighbors were assigned to each dog or to each human with respect to *MSZ* values. This process was implemented in Python using scikit-learn.

### Fitting the epigenetic age transfer function

The nearest neighbor analysis was fit using non-linear regression with the SciPy package in Python. The model fit was specified using the following formula:

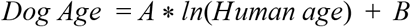

Here, “Dog age” was represented by the chronological ages of dogs, and “Human age” was the average age of the nearest human neighbors with respect to methylome similarity. The converse was performed as well, i.e. dog age was represented as the average age of the nearest dog neighbors and human age was the chronological age in humans. For the final age transfer function, the coefficients (*A*,*B*) were estimated by bootstrapping an equal number of both dogs and humans. The standard error was estimated using 1000 bootstraps.

### Mouse validation of the conserved epigenetic progression

Dog-mouse methylome similarity was calculated identically as for dog-human comparisons. A *k*-nearest neighbors analysis (as described for dogs and humans above) was repeated using the orthologous CpGs for pairwise comparisons involving mice. The mouse methylome data had a highly canalized age distribution which was different from that of the dogs or humans in our study. That is, mice had been sampled at discrete ages, we therefore visualized these data according to 0.2 year bins (**Figure 2F**).

### Identification of gene orthologs with conserved methylation trajectories

We considered 14,652 one-to-one orthologs in dogs, humans and mice that were within 2.5kb of orthologous CpGs. Among these, we identified 7,934 orthologous genes for which methylation values were available. Methylation values were then logit-transformed; multiple CpGs assigned to one gene were represented by the average methylation value. We assigned to each ortholog a ‘methylation conservation score’ using the following procedure. First, the age of each dog or mouse individual was translated to the equivalent human age using the epigenetic age translation functions built using the *k*-nearest neighbors analysis. We ranked all individuals according to their age in human years and divided this ranking into 15 quantile bins. Logit-transformed methylation values were averaged within each bin and species. For each gene and species we calculated the Spearman correlation between the gene’s methylation values and age. Genes were then ranked by *sign*(*correlation*) * − *log*_10_(*correlation*_*p value*_) within each of the three species. We computed the Euclidean norm of the three ranks and sought genes with very low norms (for which methylation was consistently among the most increasing with age across species) or with very high norms (for which methylation was consistently among the most decreasing with age across species). Significance was determined using a two-sided empirical p-value < 0.05, yielding 394 genes.

### Network analysis

We downloaded the PCNet parsimonious composite human functional interaction network from (Huang et al. 2018) and subselected gene orthologs with significantly conserved methylation trajectories (see above) resulting in a subnetwork with 355 nodes and 2003 edges. We visualized the network using Cytoscape (Shannon et al. 2003) (version 3.7) and performed community detection using clusterMaker2 (Shannon et al. 2003). To annotate modules, we performed functional enrichment using a hypergeometric test for each term within the Biological Process branch of the human Gene Ontology (GO) (Ashburner et al. 2000) and adjusted for false-discovery rate using a very strict Benjamini-Hochberg procedure (FDR < 0.001) implemented using statsmodel in python. Significant GO terms were clustered according to gene-set similarity using Enrichment Map (Merico, Isserlin, and Bader 2011), and gene modules were clustered according to their Jaccard overlap, revealing high-level functional categories (**Figure 3**).

### Developmental genes analysis

Genes were ranked according to their methylation conservation score (see above) and subdivided into 25 evenly spaced bins, separating genes with significantly conserved decreases or increases in methylation for a total of 27 bins. We then obtained PhyloP (Siepel, Pollard, and Haussler 2006) sequence conservation scores according to the orthologous CpGs assigned to each gene. Finally, we averaged the PhyloP scores in each methylation conservation score bin, estimating the 95% confidence interval by bootstrapping (**Figure S4**). We assessed the significance of the interaction between methylation conservation score and developmental gene status using ANOVA.

We restricted to orthologous CpGs profiled across dogs, humans and mice (6,906 CpGs) that were within 2.5kb of the gene bodies of all orthologous genes (‘all CpGs’). From this set, we identified CpGs near development genes (‘devCpGs’); we also controlled for the number of CpGs with 100 randomly-sampled subsets of CpGs that were equal in size from those not near developmental genes (‘not devCpGs’). We calculated the methylome similarity (as described above) based on these CpG subsets for pairwise comparisons of species (dog and human, dog and mouse). For each pairwise comparison (Species 1, Species 2), we identified the 5-nearest neighbors in Species 2 for each individual of Species 1, then binned the actual age of Species 1 into five discrete bins and calculated the average neighbor age for each bin with the 95% confidence interval estimated by bootstrapping (**Figure S5**).

### Conserved development clock analysis

We built dog and mouse epigenetic clocks with Elastic net (scikit-learn in Python) using methylation features of 439 CpGs associated with developmental gene modules (**Figure 3**). We refer to the ages predicted from this model as “epigenetic ages”. Hyperparameters were tuned using five-fold cross validation in the dog data. Performance of the final model was assessed by Spearman correlation of actual versus epigenetic age (output of the Elastic net model) for 11 dogs which had not been used for training, and in the control mice described above. For comparison, we built 100 control clocks using 100 randomly sampled sets of 439 CpGs that had been profiled in dogs and mouse but were not in developmental gene modules. For analysis involving lifespan-enhancing intervention mice, we obtained DNA methylation data profiled from whole blood from (Petkovich et al. 2017), processed as described above. We removed GHRKO from further analysis, as principal component analysis using the 439 conserved CpGs revealed clustering due to treatment. All remaining mice used in this analysis are described in **Table S3**. We applied the epigenetic clock, trained in mice or trained in dogs, and evaluated the effect of longevity-enhancing interventions using a log-likelihood ratio test.

## SUPPLEMENTARY INFORMATION

Supplementary Information accompanies this article:

Figures S1-S5

Table S1: Dog sample description

Table S2: Genes exhibiting conserved time-dependent behavior

Table S3: Mice sample description

## ACKNOWLEDGMENTS

This work is supported by the following: the California Institute for Regenerative Medicine (CIRM Epigenome Center to TI), the National Institute for Environmental Health Sciences (TI, ES014811), the National Institute of General Medical Sciences (ARC, GM108865 and TW, GM008666), the National Institute on Aging (PDA, AG031862), the National Institute of Dental and Craniofacial Research (DLB, DE022532), the Maxine Adler Endowed Chair Funds (DLB) and the Intramural Program of the National Human Genome Research Institute (AH and EAO). We also gratefully acknowledge Sabrina and Michael Mojica for providing images that document the lifespan of their beloved labrador, Cosmo.

## AUTHORS’ CONTRIBUTION

TW and TI initiated and conceptualized the study. TW carried out all experiments and implemented the main analyses. AH, EAO and DLB provided canine samples. JM, SF, BT, JFK, DLB and ARC assisted with miscellaneous analyses and provided feedback. PDA provided key input on human and animal aging. TW, TI, PDA, and EAO interpreted results and wrote the manuscript.

## AVAILABILITY OF DATA AND MATERIALS

All data and codes for analysis are available in Supplementary Materials and/or by request from the authors. Data materials availability: Sequencing data for dogs will be deposited into GEO and the accession number will be added prior to publication. The publicly available datasets used in this study can be found in the Methods section of the paper.

## REFERENCES

Alisch, Reid S., Benjamin G. Barwick, Pankaj Chopra, Leila K. Myrick, Glen A. Satten, Karen N. Conneely, and Stephen T. Warren. 2012. “Age-Associated DNA Methylation in Pediatric Populations.” Genome Research 22 (4): 623–32.

Andrews, Simon, and Others. 2010. “FastQC: A Quality Control Tool for High Throughput Sequence Data.”

Arias, Elizabeth, Melonie Heron, and Jiaquan Xu. 2017. “United States Life Tables, 2013.” National Vital Statistics Reports: From the Centers for Disease Control and Prevention, National Center for Health Statistics, National Vital Statistics System 66 (3): 1–64.

Aryee, Martin J., Andrew E. Jaffe, Hector Corrada-Bravo, Christine Ladd-Acosta, Andrew P. Feinberg, Kasper D. Hansen, and Rafael A. Irizarry. 2014. “Minfi: A Flexible and Comprehensive Bioconductor Package for the Analysis of Infinium DNA Methylation Microarrays.” Bioinformatics 30 (10): 1363–69.

Ashburner, M., C. A. Ball, J. A. Blake, D. Botstein, H. Butler, J. M. Cherry, A. P. Davis, et al. 2000. “Gene Ontology: Tool for the Unification of Biology. The Gene Ontology Consortium.” Nature Genetics 25 (1): 25–29.

Bartges, Joe, Beth Boynton, Amy Hoyumpa Vogt, Eliza Krauter, Ken Lambrecht, Ron Svec, and Steve Thompson. 2012. “AAHA Canine Life Stage Guidelines.” Journal of the American Animal Hospital Association 48 (1): 1–11.

Bell, G. I., J. H. Karam, and W. J. Rutter. 1981. “Polymorphic DNA Region Adjacent to the 5’ End of the Human Insulin Gene.” Proceedings of the National Academy of Sciences of the United States of America 78 (9): 5759–63.

Bogin, Barry, and B. Holly Smith. 1996. “Evolution of the Human Life Cycle.” American Journal of Human Biology: The Official Journal of the Human Biology Council 8 (6): 703–16.

Bolstad, Benjamin Milo. 2013. “preprocessCore: A Collection of Pre-Processing Functions.” R Package Version 1 (0).

Chen, B., C. Chen, and W. H. Hsu. 2015. “Face Recognition and Retrieval Using Cross-Age Reference Coding With Cross-Age Celebrity Dataset.” IEEE Transactions on Multimedia 17 (6): 804–15.

Cia, U. 2013. “The World Factbook 2013-14.” Central Intelligence Agency.

Ciccarone, Fabio, Stefano Tagliatesta, Paola Caiafa, and Michele Zampieri. 2018. “DNA Methylation Dynamics in Aging: How Far Are We from Understanding the Mechanisms?” Mechanisms of Ageing and Development 174 (September): 3–17.

Davis, Brian W., and Elaine A. Ostrander. 2014. “Domestic Dogs and Cancer Research: A Breed-Based Genomics Approach.” ILAR Journal / National Research Council, Institute of Laboratory Animal Resources 55 (1): 59–68.

Dreger, Dayna L., Maud Rimbault, Brian W. Davis, Adrienne Bhatnagar, Heidi G. Parker, and Elaine A. Ostrander. 2016. “Whole-Genome Sequence, SNP Chips and Pedigree Structure: Building Demographic Profiles in Domestic Dog Breeds to Optimize Genetic-Trait Mapping.” Disease Models & Mechanisms 9 (12): 1445–60.

Field, Adam E., Neil A. Robertson, Tina Wang, Aaron Havas, Trey Ideker, and Peter D. Adams. 2018. “DNA Methylation Clocks in Aging: Categories, Causes, and Consequences.” Molecular Cell 71 (6): 882–95.

Fleming, J. M., K. E. Creevy, and D. E. L. Promislow. 2011. “Mortality in North American Dogs from 1984 to 2004: An Investigation into Age-, Size-, and Breed-Related Causes of Death.” Journal of Veterinary Internal Medicine / American College of Veterinary Internal Medicine 25 (2): 187–98.

Gilmore, Keiva M., and Kimberly A. Greer. 2015. “Why Is the Dog an Ideal Model for Aging Research?” Experimental Gerontology 71 (November): 14–20.

Greeley, E. H., J. M. Ballam, J. M. Harrison, R. D. Kealy, D. F. Lawler, and M. Segre. 2001. “The Influence of Age and Gender on the Immune System: A Longitudinal Study in Labrador Retriever Dogs.” Veterinary Immunology and Immunopathology. https://doi.org/10.1016/s0165-2427(01)00336-1.

Gross, A. M., P. A. Jaeger, J. F. Kreisberg, K. Licon, K. L. Jepsen, M. Khosroheidari, B. M. Morsey, et al. 2016. “Methylome-Wide Analysis of Chronic HIV Infection Reveals Five-Year Increase in Biological Age and Epigenetic Targeting of HLA.” Molecular Cell 62 (2): 157–68.

Hannum, G., J. Guinney, L. Zhao, L. Zhang, G. Hughes, S. Sadda, B. Klotzle, et al. 2013. “Genome-Wide Methylation Profiles Reveal Quantitative Views of Human Aging Rates.” Molecular Cell 49 (2): 359–67.

Hastie, Trevor, Robert Tibshirani, Balasubramanian Narasimhan, and Gilbert Chu. 2011. “Impute: Impute: Imputation for Microarray Data.” R package version.

Horvath, S. 2013. “DNA Methylation Age of Human Tissues and Cell Types.” Genome Biology 14 (10): R115.

Horvath, Steve, and Kenneth Raj. 2018. “DNA Methylation-Based Biomarkers and the Epigenetic Clock Theory of Ageing.” Nature Reviews. Genetics 19 (6): 371–84.

Huang, Justin K., Daniel E. Carlin, Michael Ku Yu, Wei Zhang, Jason F. Kreisberg, Pablo Tamayo, and Trey Ideker. 2018. “Systematic Evaluation of Molecular Networks for Discovery of Disease Genes.” Cell Systems 6 (4): 484–95.e5.

Inoue, Mai, A. Hasegawa, Y. Hosoi, and K. Sugiura. 2015. “A Current Life Table and Causes of Death for Insured Dogs in Japan.” Preventive Veterinary Medicine 120 (2): 210–18.

Jaffe, Andrew E., and Rafael A. Irizarry. 2014. “Accounting for Cellular Heterogeneity Is Critical in Epigenome-Wide Association Studies.” Genome Biology 15 (2): R31.

Jones, Eric, Travis Oliphant, Pearu Peterson, and Others. 2015. “SciPy: Open Source Scientific Tools for Python, 2001.” URL Http://www.Scipy.Org 73: 86.

Kaeberlein, Matt, Kate E. Creevy, and Daniel E. L. Promislow. 2016. “The Dog Aging Project: Translational Geroscience in Companion Animals.” Mammalian Genome: Official Journal of the International Mammalian Genome Society 27 (7-8): 279–88.

Kowald, Axel, and Thomas B. L. Kirkwood. 2016. “Can Aging Be Programmed? A Critical Literature Review.” Aging Cell 15 (6): 986–98.

Krueger, Felix. 2015. “Trim Galore.” A Wrapper Tool around Cutadapt and FastQC to Consistently Apply Quality and Adapter Trimming to FastQ Files.

Krueger, Felix, and Simon R. Andrews. 2011. “Bismark: A Flexible Aligner and Methylation Caller for Bisulfite-Seq Applications.” Bioinformatics 27 (11): 1571–72.

Langmead, Ben, Cole Trapnell, Mihai Pop, and Steven L. Salzberg. 2009. “Ultrafast and Memory-Efficient Alignment of Short DNA Sequences to the Human Genome.” Genome Biology 10 (3): R25.

Lebeau, A. 1953. “L’âge Du Chien et Celui de L’homme. Essai de Statistique Sur La Mortalité Canine.” Bulletin de l’Academie Veterinaire de France 26: 229–32.

Li, Heng, Bob Handsaker, Alec Wysoker, Tim Fennell, Jue Ruan, Nils Homer, Gabor Marth, Goncalo Abecasis, Richard Durbin, and 1000 Genome Project Data Processing Subgroup. 2009. “The Sequence Alignment/Map Format and SAMtools.” Bioinformatics 25 (16): 2078–79.

Lowe, Robert, Carl Barton, Christopher A. Jenkins, Christina Ernst, Oliver Forman, Denise S. Fernandez-Twinn, Christoph Bock, et al. 2018. “Ageing-Associated DNA Methylation Dynamics Are a Molecular Readout of Lifespan Variation among Mammalian Species.” Genome Biology 19 (1): 22.

Madar, Aviv, Alex Greenfield, Eric Vanden-Eijnden, and Richard Bonneau. 2010. “DREAM3: Network Inference Using Dynamic Context Likelihood of Relatedness and the Inferelator.” PloS One 5 (3): e9803.

Maegawa, Shinji, George Hinkal, Hyun Soo Kim, Lanlan Shen, Li Zhang, Jiexin Zhang, Nianxiang Zhang, Shoudan Liang, Lawrence A. Donehower, and Jean-Pierre J. Issa. 2010. “Widespread and Tissue Specific Age-Related DNA Methylation Changes in Mice.” Genome Research 20 (3): 332–40.

Maegawa, Shinji, Yue Lu, Tomomitsu Tahara, Justin T. Lee, Jozef Madzo, Shoudan Liang, Jaroslav Jelinek, Ricki J. Colman, and Jean-Pierre J. Issa. 2017. “Caloric Restriction Delays Age-Related Methylation Drift.” Nature Communications 8 (1): 539.

Merico, Daniele, Ruth Isserlin, and Gary D. Bader. 2011. “Visualizing Gene-Set Enrichment Results Using the Cytoscape Plug-in Enrichment Map.” Methods in Molecular Biology. https://doi.org/10.1007/978-1-61779-276-2_12.

Miller, R. A., and N. L. Nadon. 2000. “Principles of Animal Use for Gerontological Research.” The Journals of Gerontology. Series A, Biological Sciences and Medical Sciences 55 (3): B117–23.

Ostrander, Elaine A., Robert K. Wayne, Adam H. Freedman, and Brian W. Davis. 2017. “Demographic History, Selection and Functional Diversity of the Canine Genome.” Nature Reviews. Genetics 18 (12): 705–20.

Pedregosa, Fabian, Gaël Varoquaux, Alexandre Gramfort, Vincent Michel, Bertrand Thirion, Olivier Grisel, Mathieu Blondel, et al. 2011. “Scikit-Learn: Machine Learning in Python.” Journal of Machine Learning Research: JMLR 12 (November): 2825–30.

Petkovich, Daniel A., Dmitriy I. Podolskiy, Alexei V. Lobanov, Sang-Goo Lee, Richard A. Miller, and Vadim N. Gladyshev. 2017. “Using DNA Methylation Profiling to Evaluate Biological Age and Longevity Interventions.” Cell Metabolism 25 (4): 954–60.e6.

Quinlan, Aaron R., and Ira M. Hall. 2010. “BEDTools: A Flexible Suite of Utilities for Comparing Genomic Features.” Bioinformatics 26 (6): 841–42.

Rakyan, Vardhman K., Thomas A. Down, Siarhei Maslau, Toby Andrew, Tsun-Po Yang, Huriya Beyan, Pamela Whittaker, et al. 2010. “Human Aging-Associated DNA Hypermethylation Occurs Preferentially at Bivalent Chromatin Domains.” Genome Research 20 (4): 434–39.

R Core Team. 2018. “R: A Language and Environment for Statistical Computing.” Vienna, Austria: R Foundation for Statistical Computing. https://www.R-project.org.

Rosenbloom, Kate R., Joel Armstrong, Galt P. Barber, Jonathan Casper, Hiram Clawson, Mark Diekhans, Timothy R. Dreszer, et al. 2015. “The UCSC Genome Browser Database: 2015 Update.” Nucleic Acids Research 43 (Database issue): D670–81.

Ryan, D. 2017. “MethylDackel: A (mostly) Universal Methylation Extractor for BS-Seq Experiments, 2017.”

Shannon, Paul, Andrew Markiel, Owen Ozier, Nitin S. Baliga, Jonathan T. Wang, Daniel Ramage, Nada Amin, Benno Schwikowski, and Trey Ideker. 2003. “Cytoscape: A Software Environment for Integrated Models of Biomolecular Interaction Networks.” Genome Research 13 (11): 2498–2504.

Sheffield, Nathan C., and Christoph Bock. 2016. “LOLA: Enrichment Analysis for Genomic Region Sets and Regulatory Elements in R and Bioconductor.” Bioinformatics 32 (4): 587–89.

Siepel, Adam, Katherine S. Pollard, and David Haussler. 2006. “New Methods for Detecting Lineage-Specific Selection.” In Research in Computational Molecular Biology, 190–205. Springer Berlin Heidelberg.

Stubbs, Thomas M., Marc Jan Bonder, Anne-Katrien Stark, Felix Krueger, BI Ageing Clock Team, Ferdinand von Meyenn, Oliver Stegle, and Wolf Reik. 2017. “Multi-Tissue DNA Methylation Age Predictor in Mouse.” Genome Biology 18 (1): 68.

Teschendorff, Andrew E., Francesco Marabita, Matthias Lechner, Thomas Bartlett, Jesper Tegner, David Gomez-Cabrero, and Stephan Beck. 2013. “A Beta-Mixture Quantile Normalization Method for Correcting Probe Design Bias in Illumina Infinium 450 K DNA Methylation Data.” Bioinformatics 29 (2): 189–96.

Thompson, Michael J., Bridgett vonHoldt, Steve Horvath, and Matteo Pellegrini. 2017. “An Epigenetic Aging Clock for Dogs and Wolves.” Aging 9 (3): 1055–68.

Tools, Picard. 2015. “Picard Tools.”

Urfer, Silvan R., Tammi L. Kaeberlein, Susan Mailheau, Philip J. Bergman, Kate E. Creevy, Daniel E. L. Promislow, and Matt Kaeberlein. 2017. “A Randomized Controlled Trial to Establish Effects of Short-Term Rapamycin Treatment in 24 Middle-Aged Companion Dogs.” GeroScience 39 (2): 117–27.

Vilella, Albert J., Jessica Severin, Abel Ureta-Vidal, Li Heng, Richard Durbin, and Ewan Birney. 2009. “EnsemblCompara GeneTrees: Complete, Duplication-Aware Phylogenetic Trees in Vertebrates.” Genome Research 19 (2): 327–35.

Vonholdt, Bridgett M., John P. Pollinger, Kirk E. Lohmueller, Eunjung Han, Heidi G. Parker, Pascale Quignon, Jeremiah D. Degenhardt, et al. 2010. “Genome-Wide SNP and Haplotype Analyses Reveal a Rich History Underlying Dog Domestication.” Nature 464 (7290): 898–902.

Wang, Meng, and Bernardo Lemos. 2019. “Ribosomal DNA Harbors an Evolutionarily Conserved Clock of Biological Aging.” Genome Research 29 (3): 325–33.

Wang, Tina, Brian Tsui, Jason F. Kreisberg, Neil A. Robertson, Andrew M. Gross, Michael Ku Yu, Hannah Carter, Holly M. Brown-Borg, Peter D. Adams, and Trey Ideker. 2017. “Epigenetic Aging Signatures in Mice Livers Are Slowed by Dwarfism, Calorie Restriction and Rapamycin Treatment.” Genome Biology 18 (1): 57.

Xie, Wei, Matthew D. Schultz, Ryan Lister, Zhonggang Hou, Nisha Rajagopal, Pradipta Ray, John W. Whitaker, et al. 2013. “Epigenomic Analysis of Multilineage Differentiation of Human Embryonic Stem Cells.” Cell 153 (5): 1134–48.

Yates, Andrew, Wasiu Akanni, M. Ridwan Amode, Daniel Barrell, Konstantinos Billis, Denise Carvalho-Silva, Carla Cummins, et al. 2016. “Ensembl 2016.” Nucleic Acids Research 44 (D1): D710–16.

