## Supplementary Information: Figure S1-Figure S5 and Table S1 for "Quantitative translation of dog-to-human aging by conserved remodeling of epigenetic networks"

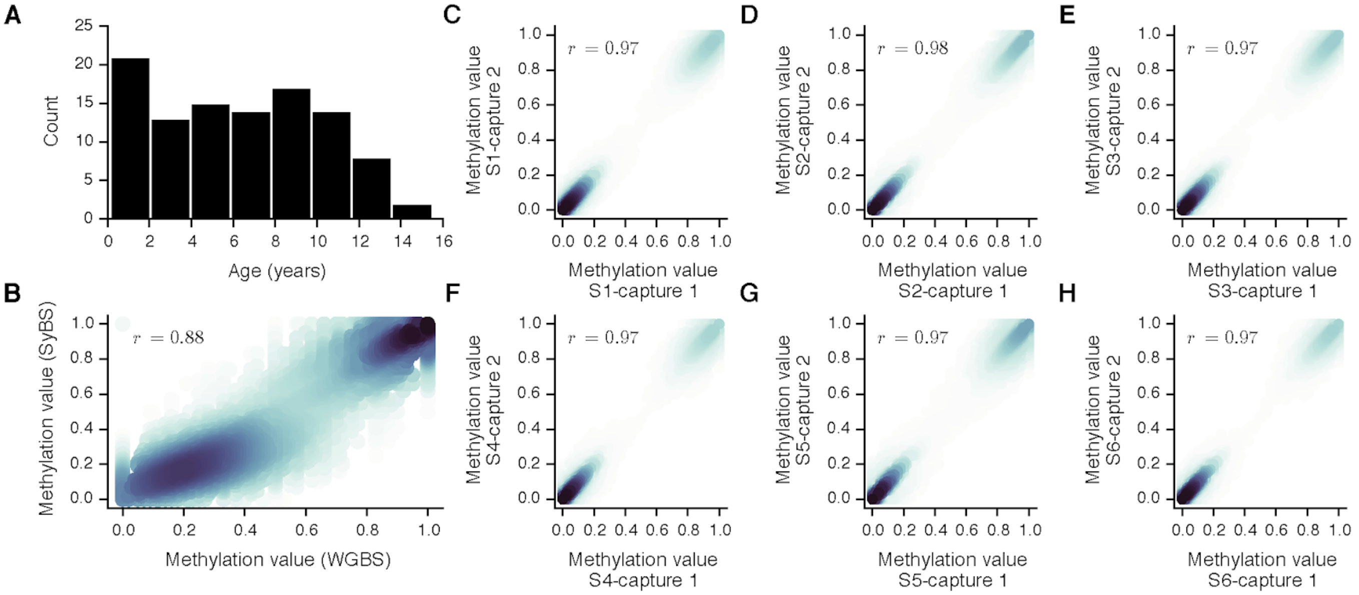


**Figure S1: Concordance of SyBS with non-targeted sequencing. (A)** The distribution of ages of 104 dogs used in this study, depicted as a histogram. (**B-F**) Concordance of SyBS values for five canine DNA samples (S2-S6), for which two independent captures were performed. The x-axis represents values obtained for the first capture, and the y-axis represents values obtained for the second capture. The Pearson correlation of the two captures is shown. Colors represent density of observations.

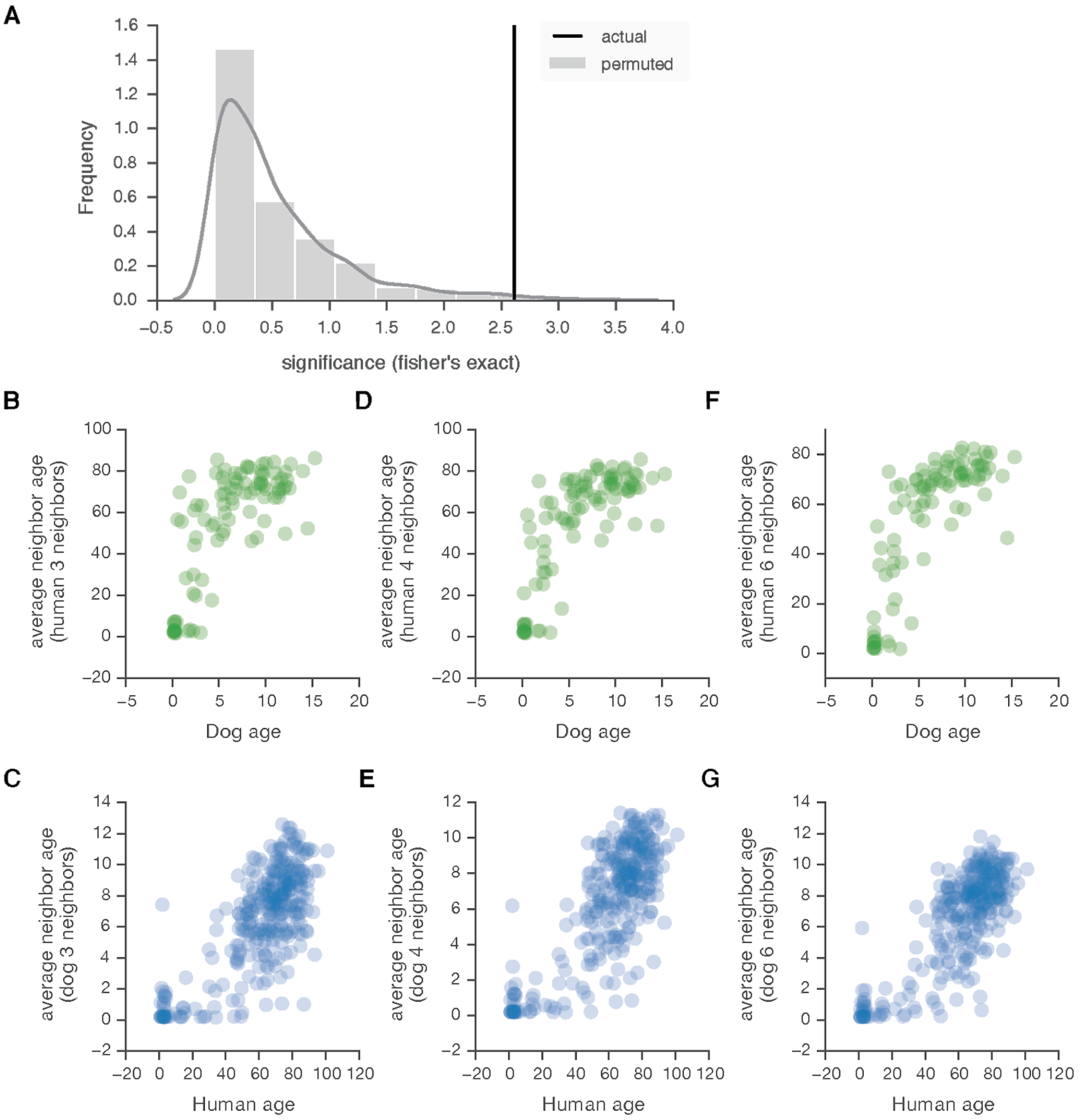


**Figure S2: Evaluating methylome similarities observed for dogs and humans.** (**A**) Black line: the observed p-value of association between methylome similarity for each pair of species and age. P-value computed by Fisher's exact test. The 'significance' ($-\log_{10}$(p-value) ) is compared to those obtained from 1000 randomizations in which dog and humans were permuted (gray bars). (**B-G**) Varying the number of nearest neighbors *k* in dog-to-human age alignments. The actual ages of dogs (in years, x-axis, **B,D,F**) or humans (x-axis, **C,E,G**) versus the average age of *k* human or dog neighbors (in years, y-axis), respectively, when varying *k*: (**B,C**) *k* = 3, (**D,E**) *k* = 4 and (**F,G**) *k* = 6. Results for *k* = 5 are shown in the main text (**Figure 2**).

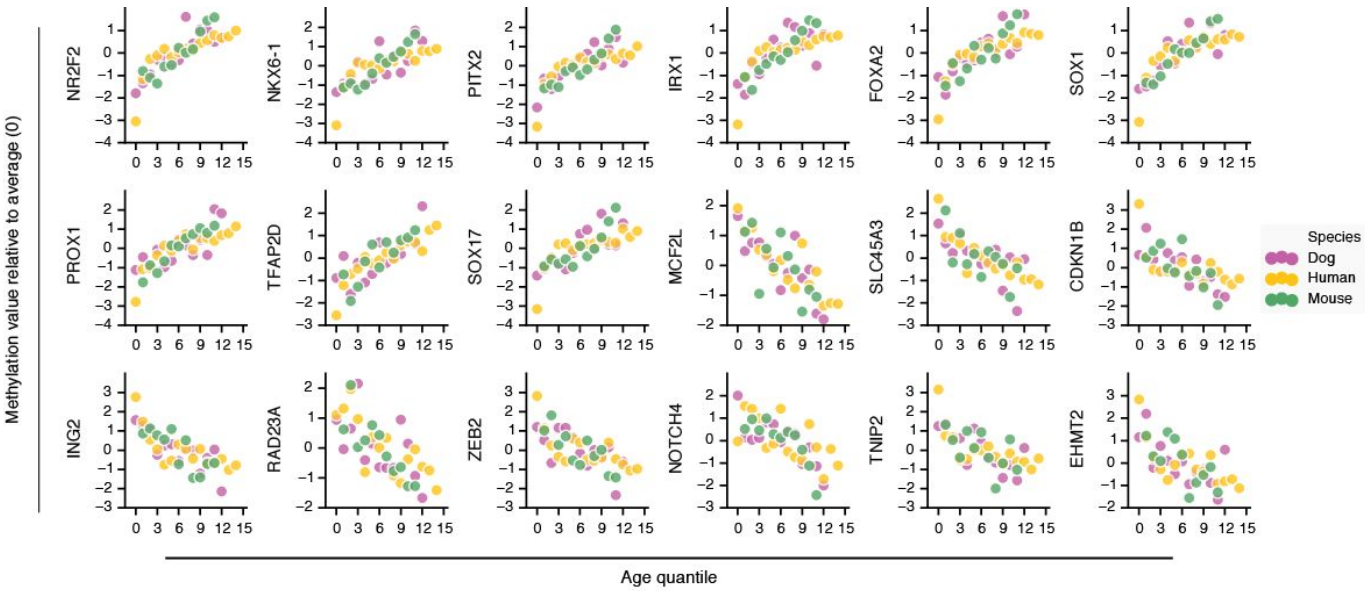


**Figure S3: Representative genes exhibiting conserved methylation changes with age.** All dog, human and mouse individuals are ranked according to their ages translated to dog years (see text), then sorted into 15 age bins (quantiles, x-axis). Eighteen out of 394 orthologs for which CpG methylation values (y-axis) exhibit conserved direction of change with age are shown. Methylation values are normalized by the mean and standard deviation within each species. Colors indicate species.

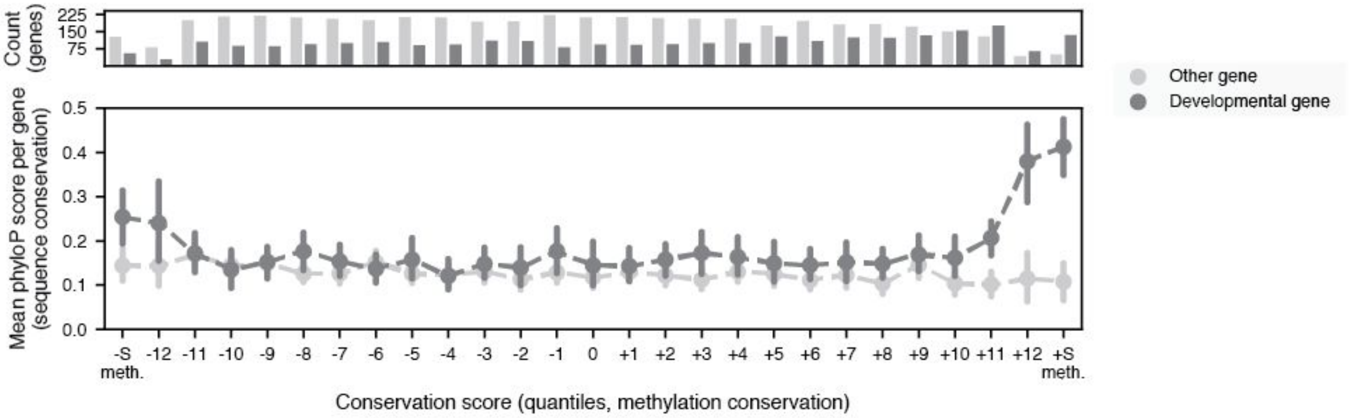


**Figure S4: Relationship between sequence constraints and conserved methylation changes with age for developmental gene modules.** We analyzed 7,942 genes with 1:1 orthologs across humans, dogs and mice and for which methylation data were available in all species. For each ortholog we computed a methylation conservation score (x-axis, binned into quantiles) and a sequence conservation score (y-axis, PhyloP). The orthologs with most significant conserved increases or decreases in methylation with age (198 and 196 genes respectively) are indicated by −S and +S at the extreme ranges of x. Vertical error bars indicate the 95% confidence interval estimated by bootstrapping. Orthologs are treated separately according to their developmental status (dark points, developmental gene; light points, non-developmental gene). The plot reveals a strong association between methylation conservation and sequence conservation, in a manner that is specific to developmental genes. The top barplot depicts the number of genes within each quantile bin. Further details on computation of conservation scores are given in **Methods**.

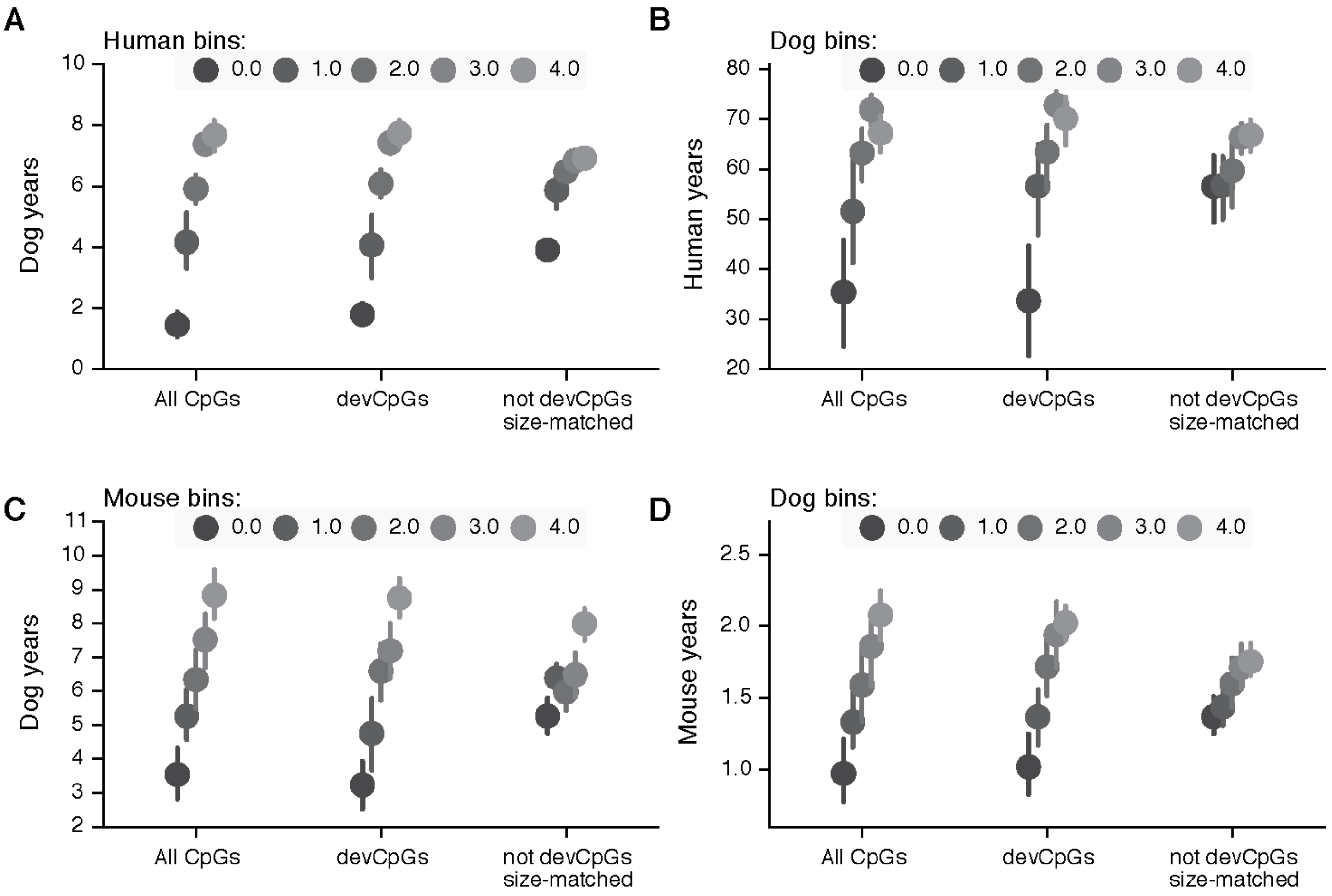


**Figure S5: Dependence of the conserved epigenetic progression on CpGs within developmental gene modules.** The plots show alignments of age across species based on methylome similarity: (**A**) dogs versus humans, (**B**) humans versus dogs, **(C)** dogs versus mice, **(D)** mice versus dogs. Within each species comparison, methylome similarity is calculated using All CpGs, CpGs near development genes (devCpGs), or randomly sampled CpG sets of equal size but not near development genes (not devCpGs size-matched). In each comparison, the x-axis represents increasing age bins for one species (quintiles), and the y-axis represents the average ages of the 5 nearest neighbors in the opposite species according to methylome similarity. Vertical bars show the 95% confidence intervals of the averages obtained from 1000 bootstrapped samplings. Note that in all comparisons (A-D), the strong trends observed for All CpGs are preserved for devCpGs but are substantially weakened for the size-matched controls.

**Table S1: Dog sample description†**

| Sample | Age (years) | Breed | Center | | Sample | Age (years) | Breed | Center |
|---|---|---|---|---|---|---|---|---|
| 2650 | 0.4 | Lab | Davis | | 2623 | 15.5 | Lab | Davis |
| 7935 | 0.5 | Lab | Davis | | 7936 | 0.5 | Lab | Davis |
| 3352 | 0.6 | Lab | Davis | | 3018 | 1.8 | Lab | Davis |
| 3591 | 1.0 | Lab | Davis | | 2519 | 2.5 | Lab | Davis |
| 5866 | 1.7 | Lab | Davis | | 2538 | 5.7 | Lab | Davis |
| 3033 | 2.2 | Lab | Davis | | 2967 | 6.3 | Lab | Davis |
| 2661 | 2.2 | Lab | Davis | | 5856 | 6.6 | Lab | Davis |
| 7008 | 2.3 | Lab | Davis | | 2667 | 7.6 | Lab | Davis |
| 7295 | 2.4 | Lab | Davis | | 2935 | 8.3 | Lab | Davis |
| 2640 | 2.5 | Lab | Davis | | 3592 | 9.3 | Lab | Davis |
| 2564 | 2.5 | Lab | Davis | | 6537 | 10.0 | Lab | Davis |
| 7007 | 2.6 | Lab | Davis | | 2624 | 10.6 | Lab | Davis |
| 2569 | 3.2 | Lab | Davis | | 2888 | 10.9 | Lab | Davis |
| 3654 | 4.3 | Lab | Davis | | 3274 | 10.9 | Lab | Davis |
| 3539 | 4.7 | Lab | Davis | | 2714 | 11.0 | Lab | Davis |
| 3209 | 4.8 | Lab | Davis | | 3696 | 11.3 | Lab | Davis |
| 2616 | 4.8 | Lab | Davis | | 2638 | 11.4 | Lab | Davis |
| 2665 | 5.4 | Lab | Davis | | 2664 | 11.4 | Lab | Davis |
| 2928 | 5.5 | Lab | Davis | | 2542 | 11.5 | Lab | Davis |
| 2554 | 5.5 | Lab | Davis | | 2266 | 11.9 | Lab | Davis |
| 2597 | 5.6 | Lab | Davis | | 2477 | 12.1 | Lab | Davis |
| 2583 | 5.6 | Lab | Davis | | 2489 | 12.3 | Lab | Davis |
| 2613 | 5.6 | Lab | Davis | | 2270 | 12.7 | Lab | Davis |
| 2557 | 5.6 | Lab | Davis | | 2849 | 14.5 | Lab | Davis |
| 2810 | 6.1 | Lab | Davis | | 3592 | 9.3 | Lab | Davis |
| 3589 | 6.3 | Lab | Davis | | 23855 | 0.1 | Lab | NHGRI |
| 2625 | 6.4 | Lab | Davis | | 22139 | 0.2 | Lab | NHGRI |
| 2649 | 6.4 | Lab | Davis | | 23848 | 0.2 | Lab | NHGRI |
| 7335 | 6.6 | Lab | Davis | | 23852 | 0.2 | Lab | NHGRI |
| 7153 | 6.7 | Lab | Davis | | 23840 | 0.2 | Lab | NHGRI |
| 6750 | 6.8 | Lab | Davis | | 23826 | 0.2 | Lab | NHGRI |
| 3054 | 6.9 | Lab | Davis | | 23820 | 0.2 | Lab | NHGRI |
| 2619 | 7.3 | Lab | Davis | | 23824 | 0.2 | Lab | NHGRI |
| 7046 | 7.8 | Lab | Davis | | 23792 | 0.2 | Lab | NHGRI |
| 3073 | 7.8 | Lab | Davis | | 23800 | 0.2 | Lab | NHGRI |
| 3436 | 8.0 | Lab | Davis | | 23796 | 0.2 | Lab | NHGRI |
| 3255 | 8.0 | Lab | Davis | | 25293 | 1.8 | Lab | NHGRI |
| 2500 | 8.2 | Lab | Davis | | 9137 | 3.0 | Lab | NHGRI |
| 3220 | 8.3 | Lab | Davis | | 31279 | 3.1 | Lab | NHGRI |
| 7328 | 8.3 | Lab | Davis | | 25415 | 3.8 | Lab | NHGRI |
| 2604 | 8.4 | Lab | Davis | | 9127 | 5.3 | Lab | NHGRI |
| 2668 | 8.5 | Lab | Davis | | 22211 | 9.5 | Lab | NHGRI |
| 2776 | 8.8 | Lab | Davis | | 31117 | 9.5 | Lab | NHGRI |
| 2927 | 9.1 | Lab | Davis | | 30379 | 11.7 | Lab | NHGRI |
| 2755 | 9.2 | Lab | Davis | | 7725 | 12.6 | Lab | NHGRI |
| 3328 | 9.5 | Lab | Davis | | 13295 | 13.3 | Lab | NHGRI |
| 2862 | 9.6 | Lab | Davis | | D1 | 1.5 | Mix | CV |
| 2903 | 9.7 | Lab | Davis | | D6 | 1.9 | Mix | CV |
| 2916 | 9.8 | Lab | Davis | | D4 | 4.2 | Mix | CV |
| 3151 | 9.9 | Lab | Davis | | D5 | 4.6 | Mix | CV |
| 6529 | 9.9 | Lab | Davis | | D2 | 6.8 | Pug | CV |
| 2747 | 10.0 | Lab | Davis | | | | | |

**†** The table lists the 104 dogs used in this study: the sample ID, age in years and center where the sample was obtained are shown. Abbreviations: Labrador retriever (Lab), Mixed breed (Mix), UC Davis (Davis), National Human Genome Research Institute (NHGRI), Consented Volunteer (CV).
